# Detection of probable neuronal gene expression changes in skin biopsies from patients with paclitaxel-induced peripheral neuropathy

**DOI:** 10.1101/2025.10.15.682581

**Authors:** Andy Wangzhou, Surendra Dasari, Victoria Pastor, Ryan R. Ju, Michelle Mauermann, Sybil Hrstka, Sandra Rieger, Diana Tavares-Ferreira, Nathan P. Staff, Theodore J. Price

## Abstract

Our inability to obtain nerve samples from the vast majority of neuropathic pain patients impedes our ability to understand the disease, creates challenges in understanding mechanisms in specific patient populations, and limits our ability to make treatment decisions based on quantifiable molecular data. Fields like oncology have overcome these problems to take advantage of the insight that sequencing offers for understanding mechanisms of disease and have leveraged these molecular insights to dramatically change the treatment landscape in the past decade. Here we tested the hypothesis that skin biopsies could be used to gain insight into neuronal transcriptomic changes in patients with paclitaxel-induced peripheral neuropathy (PIPN). Our analysis reveals that hundreds of differentially expressed genes (DEGs) found through bulk RNA sequencing in these skin biopsies are likely contributed by dorsal root ganglion (DRG) neuronal axons and/or terminals. Up-regulated genes were representative of broad class of nociceptors whereas down-regulated genes were associated with putative injured DRG neurons expressing the *PDIA2* gene. DEGs that could be confidently associated with specific subsets of skin cells were mostly expressed by keratinocytes supporting a growing literature tying keratinocyte-neuron communication abnormalities to pain in PIPN. We validated these findings by analyzing additional previously published datasets and through prospectively conducted spatial transcriptomics experiments on human skin biopsies. Our findings warrant further assessment of skin biopsies in additional neuropathic pain populations to gain insight into DRG neuron changes that have previously been thought to be inaccessible in routine clinical or scientific assessments in most patients.

## Introduction

Peripheral neuropathy caused by chemotherapeutic treatment is an important clinical problem that is increasingly frequent as cancer survivorship continues to improve [19; 33; 34; 37; 45]. Paclitaxel-induced peripheral neuropathy (PIPN) is a subset of chemotherapy-induced peripheral neuropathy (CIPN) and is particularly frequent and a dose-limiting side effect of this mainstay of treatment for several cancers, like breast cancer. There are no approved treatments for PIPN, but animal models of PIPN, and CIPN in general, have revealed mechanisms that may explain why this type of neuropathy occurs, and why it often develops into a peripheral neuropathy that persists even after paclitaxel treatment has ended [36]. These mechanisms include die-back of axonal endings from the skin, mitochondrial dysfunction and increased oxidative stress within the dorsal root ganglion (DRG) and peripheral nerves [36; 45], and changes in gene expression that cause DRG neurons to become hyperexcitable causing neuropathic pain [16; 20]. Validating these mechanisms in humans is essential to inform appropriate treatment development.

While there have been many advances in understanding how the human peripheral nervous system changes in neuropathic pain, these methods mostly rely on recovery of tissues from organ donors [35], or from rare surgeries where the DRG is excised, like thoracic vertebrectomy and C1/C2 fusion surgeries [1; 26; 29]. These procedures cannot be done in almost all PIPN patients. Skin biopsies offer an accessible alternative. DRG neurons contain long axons that innervate every part of the body, and the skin is one of the most densely innervated organs. mRNAs are transported into these axons in all species where this has been examined [25], and the local translation of these mRNAs has been shown to contribute to neuropathic pain and increased excitability of nerve endings [10; 12; 21; 22; 31; 43]. In diabetic peripheral neuropathy patients, the localization of these mRNAs appears to change in peripheral nerves, suggesting this may also occur in other human neuropathies [40]. The goal of our study was to test whether skin biopsies of patients with PIPN would display altered mRNA expression profiles consistent with changes in axonal mRNA distribution.

Here we used distal leg skin biopsies from female individuals with PIPN and from matched age and sex control individuals with no history of neuropathy. Intraepidermal nerve fiber (IENF) densities were measured in these skin biopsies, and no significant differences were detected between the PIPN and control groups (Table 1) [38]. This is important because neuronal mRNAs can only be measured if the nerve endings are still present in the skin biopsy samples. Furthermore, this mRNA data potentially adds diagnostic value in symptomatic patients with normal IENF density [5]. We assembled additional skin biopsy datasets to examine whether these neuronal genes can be detected across different patient groups, finding robust changes in neuronally-enriched genes in PIPN and across other biopsy cohorts. Our findings support the conclusion that skin biopsies can provide a “window” into DRG neuronal gene expression changes in peripheral neuropathy patients. pointing toward a new diagnostic paradigm accessible in most neuropathic pain patients who do not have excessive axonal die-back from the biopsied area.

**Table 1.** Patient demographics and neuropathy measures. Sex and age are shown for three healthy controls and three patients with chemotherapy-induced peripheral neuropathy (CIPN) following paclitaxel treatment. Skin biopsies were collected at the indicated time points after CIPN diagnosis. QLQ-CIPN20 scores are presented as the total score, sensory-specific or motor-specific scores. Neurological impairment score of the lower limbs (NIS-LL) and IENF are also included.

| Study ID | Treatment | Sex | Age at Consent | Weeks from Neuropathy to Consent | NIS-LL | CIPN20 Score<br>(Total: 19-76) | CIPN20 Score<br>(Sensory: 9-36) | CIPN20 Score<br>(Sensory: 3-12) | IENF |
| --- | --- | --- | --- | --- | --- | --- | --- | --- | --- |
| CIPN001 | Pctx | F | 70 | 35 | 2 | 31 | 22 | 3 | 8.9 |
| CIPN002 | Pctx | F | 70 | 31.43 | 12 | 34 | 22 | 4 | 5.3 |
| CIPN003 | Pctx | F | 60 | 4.86 | 4 | 25 | 16 | 3 | 9.6 |
| CTRL005 | CTRL | F | 70 | n/a | 0 | 19 | 9 | 3 | 14.9 |
| CTRL006 | CTRL | F | 65 | n/a | 0 | 19 | 9 | 3 | 7.6 |
| CTRL007 | CTRL | F | 64 | n/a | 0 | 19 | 9 | 3 | 7.6 |
| **Average (Pctx vs. CTRL)** |  |  | **67/66** | **23.76** | **6** | **30/19** | **20/9** | **3.3/3.0** | **7.9/10.3** |

## Methods

### Patient Cohort

We used data from a previously-generated cohort of three patients with neuropathic dysesthesias and numbness caused by paclitaxel treatment, whom had epidermal nerve fiber density processed through standard clinical laboratory practice[7], which was within normal limits and equivalent to three age- and sex-matched controls. The patient cohort and data generation are described in [38]. The patient information and neuropathy scores including other metadata are presented in Table 1 (reprint from [38]). All participants were recruited, interviewed, and examined by a neurologist with peripheral nerve subspecialty training (NPS). Additional information of the clinical procedures and nerve fiber staining can be found in the Supplement File 1.

### Skin processing and RNA sequencing

We reanalyzed data generated in Staff et al., for the experiments described here. Details on sample generation and RNA sequencing were described in detail in that previous publication [38]. Briefly, skin biopsies were collected from distal limb sites, followed by library preparation and RNA-seq. Because of constraints in the downstream annotation pipeline, only protein-coding genes were retained for analysis. Protein-coding gene annotation was based on GENCODE v32, which is the version underlying the widely used 10x Genomics reference (2020-A). Differential gene expression analysis was performed using DESeq2 (v1.34.0). Compared to the original publication, which applied a minimum count threshold of 20 to maximize sensitivity for detecting broad transcriptional differences, we increased this threshold to 50. This more stringent filtering was employed to minimize the contribution of ultralow-expressing genes that could confound the neuronal gene signal identified in skin, thereby increasing confidence that detected neuronal gene expression reflects a signal potentially originating from neuronal cells rather than genes natively expressed at low levels in skin tissue.

### Neuronal gene enrichment analysis

To isolate the differentially expressed genes (DEGs) that were likely contributed by nerve endings and axons in skin tissue, we compared multiple single-nucleus RNA sequencing (snRNA-seq) and single-cell RNA-sequencing (scRNA-seq) datasets with the original bulk RNA-seq data to isolate this signal. An overview of this workflow is presented in Fig. 1. First, we used a previously published skin scRNA-seq atlas to identify transcripts likely originating from skin-resident cell types. This atlas comprises 273,178 skin cells aggregated from 14 datasets, representing a total of 85 individuals. This dataset includes skin samples from 15 distinct anatomical regions, encompassing normal skin tissues obtained from foreskin circumcisions, reduction abdominoplasties or mammoplasties, skin grafts, forearm and leg biopsies, palm, sole, hip, facial, inguinoiliac region, and scalp excisions, etc. [30]. Clustering and cell-type annotations were supplied by the authors [30]. Genes detected in > 10% cells within any given cell-type were considered expressed in that cell-type. DEGs not detected in any skin cell-types were subsequently examined for expression in sensory neurons from human DRG. For this analysis, we used a recently published scRNA-seq atlas of 132 human DRGs and deep single-soma RNA-seq dataset [3; 4]. The same >10% cell detection threshold was applied. DEGs identified in either skin cell types or sensory neurons were then evaluated for cell-type specificity (restricted vs. pan-cellular expression), followed by pathway and gene ontology (GO) analyses using Enrichr [13; 48].

**Figure 1.**
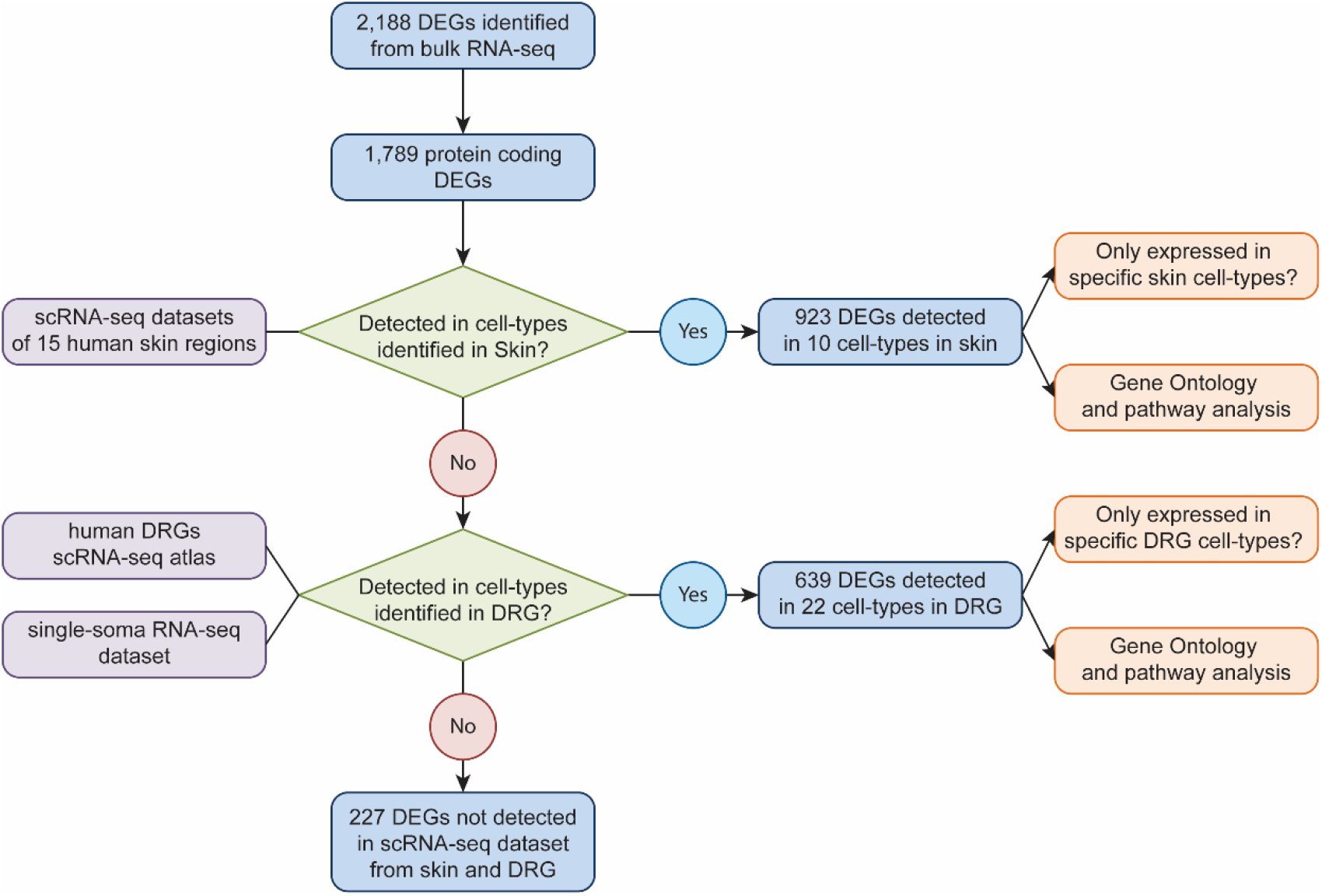
Experimental workflow. A flowchart showing the procedure used to identify genes associated with specific skin cell types, and specific DRG nociceptor subtypes detected in nerve endings of skin biopsy samples.

### Validation in other skin biopsy datasets

To evaluate whether the detection of presumed neuronal transcripts generalized across additional neuropathic skin biopsy datasets, we applied the same analytical workflow used for the PIPN cohort to three independent bulk RNA-seq datasets: complex regional pain syndrome (CRPS), diabetic painful neuropathy (DPN), and psoriatic arthritis skin biopsies[9; 18; 44]. Protein-coding genes detected in each dataset were compared against the same human skin scRNA-seq reference atlas and human DRG neuronal transcriptomic datasets described above. Genes detected in >10% of cells within a given skin cell type or neuronal subtype were considered expressed in that population. The fraction of expressed genes attributable to skin cell types versus presumed neuronal populations was then compared between disease and control conditions within each dataset.

### Xenium

Xenium In Situ Gene Expression protocols were performed according to the manufacturer’s instructions from 10x Genomics. Briefly, skin tissue was formalin-fixed paraffin-embedded as previously described in [7], and was then sectioned at 5 µm and mounted onto Xenium slides. Tissues sections were deparaffinized, dehydrated, decrosslinked, before overnight probe hybridization. Cell segmentation staining was also performed using the Xenium Cell Segmentation Staining add-on according to the manufacturer’s instructions. Following tissue preparation and staining, slides were loaded onto the Xenium Analyzer, where the remaining in situ chemistry, imaging, and transcript decoding were performed using the Xenium analysis workflow.

Six tissue sections from three PIPN donors (CIPN001, CIPN002, and CIPN003) were profiled. Transcript detections were filtered to include annotated genes with quality values ≥20, retaining high-confidence transcript calls. A curated list of 639 axon-associated genes (Suppl. Table 1) was cross-referenced with Xenium transcript output files, identifying 36 detected genes. Genes were grouped into seven functional categories: neurofilament/cytoskeletal, ion channel, axon guidance/adhesion, synaptic/vesicular, trophic/signaling, myelination/Schwann cell, and transcription/RNA binding. Using Python (v3.12.12), transcript counts were extracted from transcripts.parquet output files, aggregated by donor and section, and visualized as donor-stratified stacked bar plots. Spatial distributions of selected transcripts (*NEFH, TUBB3*, and *SCN9A*), were overlaid on tissue morphology images (DAPI and alphaSMA/Vimentin - interior protein) to visualize their localization within skin sections and zoomed views of selected transcript markers were generated in 10x Genomics Xenium Explorer v4.1.0.

### Immunohistochemistry - PGP9.5

PGP9.5 immunohistochemistry was performed on FFPE skin sections. Briefly, sections were deparaffinized, rehydrated, subjected to antigen retrieval, blocked, and incubated with anti-PGP9.5 primary antibody (ab108986, Abcam) followed by fluorescent secondary antibody. Sections were counterstained with DAPI and imaged on an Olympus FV4000 confocal microscope.

## Results

In our reanalysis of previously published data on bulk RNA sequencing from distal leg skin biopsies of patients suffering from PIPN, but with intact intraepidermal nerve fibers versus control skin biopsies, we found a total of 2,188 DEGs using a threshold of FDR less than 0.05 and a log2 fold change of greater than 0.5. Of these genes, 1,553 were up-regulated while 625 were down-regulated. Among this total number of DEGs, 1,789 were protein coding genes and among these 1,226 were up-regulated and 563 were down-regulated (Suppl. Table 1). Our remaining analysis was aimed at understanding how many of these DEGs could have been contributed by neuronal axons within the skin.

We first utilized scRNA-seq datasets from human skin tissues [30] with the goal of eliminating transcripts from the DEGs that could be confidently called as contributed by skin cells. Of these, we identified 923 out of 1,789 protein coding DEGs detected in 10 major cell-types from human skin samples (Fig 2A, Suppl. Table 1). Among these 923 genes, 400 of them showed decreased expression in PIPN patients. GO enrichment analysis shows that these genes are mainly involved in negative regulation of angiogenesis, and MHC II antigen presentation (Fig. 2B). This finding is consistent with the known mechanism of action of many chemotherapeutic drugs that target angiogenesis-related pathways. Transcription factor analysis on these down-regulated DEGs points out *TCF21*, which is known for inhibition of tumor-associated angiogenesis through PI3K and ERK signaling, and modulate fibroblast activation after injury [2; 6].

**Figure 2.**
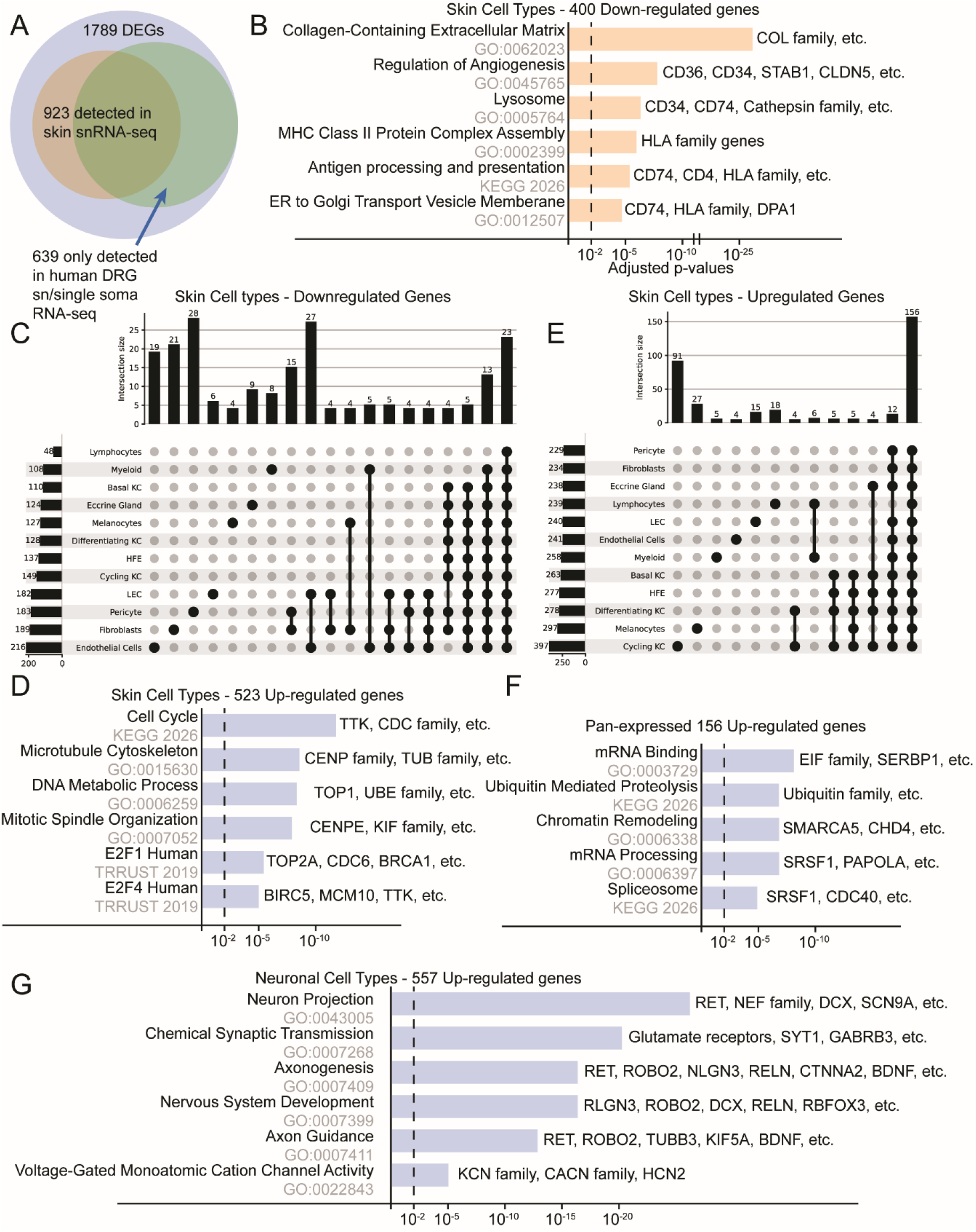
DEGs identified in skin cell-types. A) Proportion of DEGs identified in snRNA-seq dataset from skin and DRGs. B) GO and pathway analysis of 402 down-regulated DEGs potentially from skin cell-types, and C) their cell-type specificity is presented with the upset plot. The vertical bar shows the number of genes detected in the cell-type or cell-type combination represented by the dots below. The horizontal bar represents the number of genes detected in the corresponding cell-type. D) GO and pathway analysis of 524 up-regulated DEGs potentially from skin cell-types, and E) their cell-type specificity. F) GO and pathway analysis of 156 up-regulated DEGs detected in all skin cell-types. G) GO and pathway analysis for 557 up-regulated DEGs potentially from nerve endings in the skin.

Pathway analysis also points out transforming growth factor (TGF) -beta regulation of the extracellular matrix (ECM) as a key downregulated signaling pathway in the skin cells among downregulated genes. Although many of these downregulated genes were expressed across cell types, the most enriched was endothelial cells with 54 of the down-regulated genes coming solely from this cell type. The next most common cell types in this analysis were fibroblasts with 31 downregulated DEGs and myofibroblasts with 19 downregulated DEGs (Fig. 2C).

We also identified 523 genes with increased expression levels in the DEG list that we could confidently call as expressed in skin cells (Suppl. Table 1). GO enrichment analysis on these 523 genes showed enrichment in processes involving microtubule dynamics and nuclear functions aligning with the basic characteristics of cycling keratinocytes. Pathway and GO analysis pointed to cell cycle and E2F family transcription factors as upregulated pathways in these cells (Fig. 2D). These pathways and GO terms enrichments were driven primarily by the 94 genes exclusively detected in cycling keratinocytes (Fig. 2E), which was the most abundant cell type with enrichment of DEGs that were upregulated. The findings suggest that this cell type is involved in an active process in the skin of patients with PIPN. We next examined the 156 genes shared across all skin cell types. Pathway and GO analyses showed enrichment for mRNA-processing functions, suggesting elevated post-transcriptional regulation in PIPN skin cells (Fig. 2F). Recent studies suggest that keratinocytes play a pathological role in regulating the activity of nerve endings in PIPN specifically and also more broadly in CIPN to promote pain [8; 23; 24; 39].

Out of the remaining 866 DEGs that were not detected in any cell-types identified from the skin tissues, 639 were confidently detected in neuronal cell-types from human DRGs (Fig. 2A). This finding is consistent with the notion that many of these mRNAs are found in the nerve endings and/or axons of DRG neurons that innervate the skin.

Among the 639 DEGs, 557 were up-regulated (Suppl. Table 1). These genes were mostly expressed across neuronal cell types suggesting that a large proportion of this response was a pan-neuronal reaction to chemotherapy treatment. These presumed neuronal genes include 5 voltage gated sodium channels, 17 voltage gated potassium channels, 8 glutamate receptors, and many other channels including but not limited to voltage-gated calcium channels, *TRPM3*, and others. Many of these ion channels have previously been associated with pain either in clinical studies or in animal models of CIPN [15; 20; 49]. GO analysis showed synaptic transmission functions, axonogenesis, and neuro-development related terms for this gene set (Fig 2G). An implication of this finding is that changes in neuronal gene expression or trafficking of mRNAs in axons can be measured in skin biopsies of humans with neuropathic pain.

Among the 639 DEGs, 82 were down-regulated (Suppl. Table 1). Many of these genes showed reduced expression across the cell types, including Prostaglandin D_ receptor (PTGDR), and Protein Disulfide Isomerase Family A Member (PDIA2). *PTGDR* were previously identified to be associated with neuroinflammatory responses. We have also previously identified *PDIA2* as a potential marker of axonal injury in human DRG neurons [1]. This finding supports the conclusion that injured neurons, which may have died back from their terminal endings in the skin, show de-enrichment for their axonal mRNAs in skin biopsies from PIPN patients.

To further validate that the presumed neuronal transcripts identified in the bulk RNA-seq datasets are physically present within human skin tissue, we performed Xenium spatial transcriptomic profiling on six skin biopsy sections obtained from the three individuals with PIPN (Fig. 3A). We designed the Xenium panel to include neuronal and nerve-associated transcripts and processed the samples with the Xenium Cell Segmentation Staining add-on, including DAPI and αSMA/Vimentin (interior protein), to support cell segmentation and aid visualization of nerve-associated structures (Fig. 3B). Among the presumed neuronal genes identified in the PIPN analysis, 36 were represented in the Xenium panel, and all 36 were detected across the skin biopsy sections (Fig. 3C). These included canonical neuronal markers and ion channel transcripts such as *NEFH, TUBB3*, and *SCN9A*, which displayed sparse but spatially localized expression patterns including signals near superficial epidermal regions, consistent with the expected distribution of cutaneous nerve-associated structures (Fig. 3D-E). Because the rabbit anti-PGP9.5 antibody was not compatible with the Xenium workflow, PGP9.5 immunostaining was performed on adjacent FFPE sections (Fig. 3F). These sections were used to provide independent anatomical confirmation of intraepidermal and dermal nerve fiber patterns corresponding to regions where neuronal transcripts were detected in Xenium data. These findings provide spatial validation that neuronal transcripts identified through bulk RNA sequencing can be directly detected within human skin biopsies and support the conclusion that at least a substantial subset of the inferred neuronal DEGs originate from cutaneous sensory nerve endings and axons.

**Figure 3.**
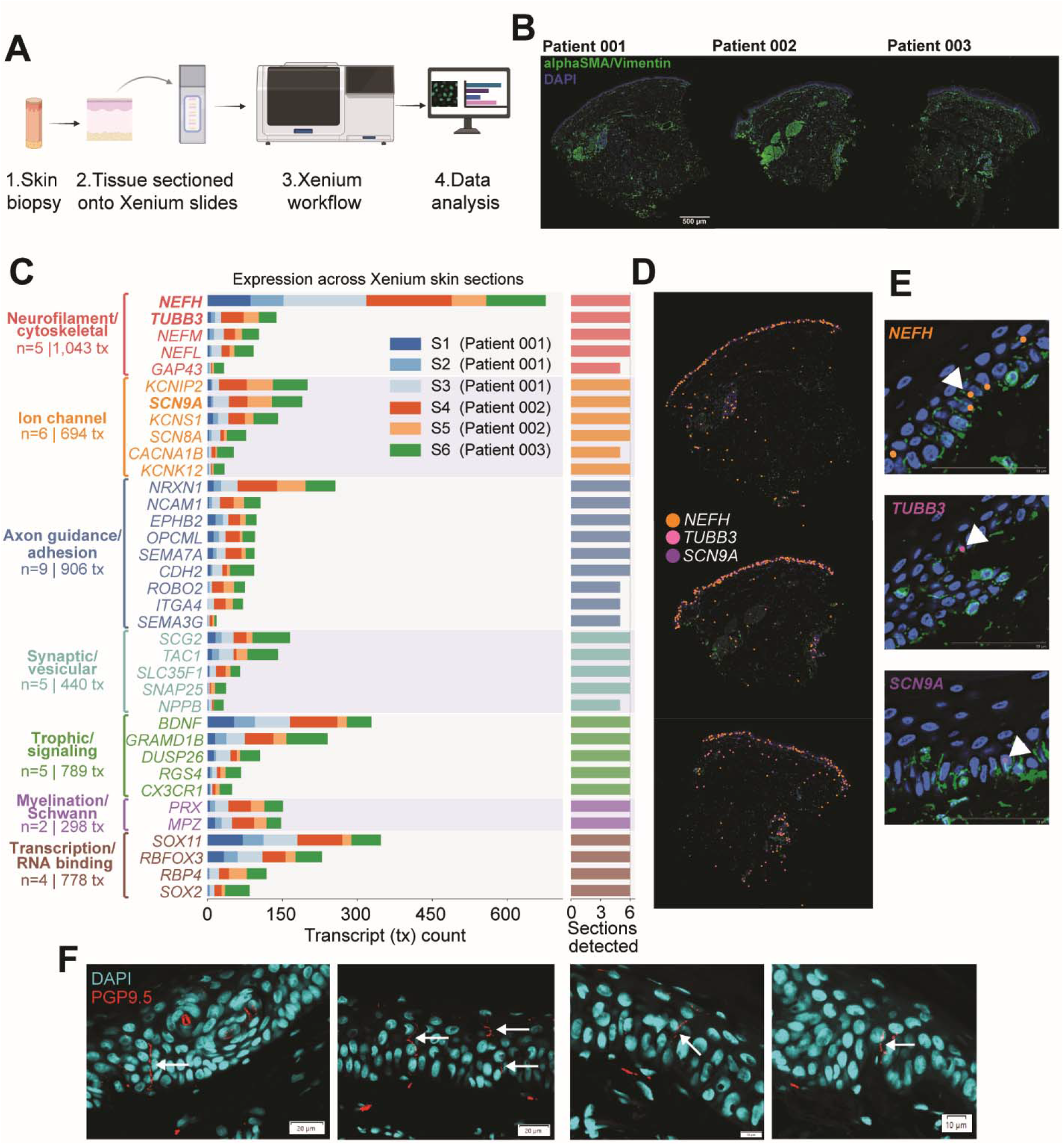
Xenium spatial transcriptomic and histological profiling of human skin biopsies identifies cutaneous nerve-associated transcripts. A) Overview of the experimental workflow. B) Representative whole-section Xenium images from three patients showing alphaSMA/vimentin (interior protein, green) and DAPI (blue), highlighting overall tissue architecture and stromal compartments. Scale bar, 500 µm. C) Expression summary of nerve-associated transcripts detected across six Xenium skin sections (three sections from Patient A, two from Patient B, one from Patient C).Genes are grouped by functional class. Bar lengths indicate transcript counts, and the adjacent right bar denotes the number of sections in which each gene was detected. D) Spatial localization of selected neuronal markers *NEFH, TUBB3*, and *SCN9A* across representative skin sections. E) Zoomed views generated in 10x Genomics Xenium Explorer (v4.1.0) of *NEFH, TUBB3*, and *SCN9A* transcripts (arrowheads). F) Immunohistochemical validation using PGP9.5 (red) with DAPI nuclear counterstain (cyan), showing sparse intraepidermal and dermal nerve fibers (arrows). Scale bars as indicated.

**Figure 4.**
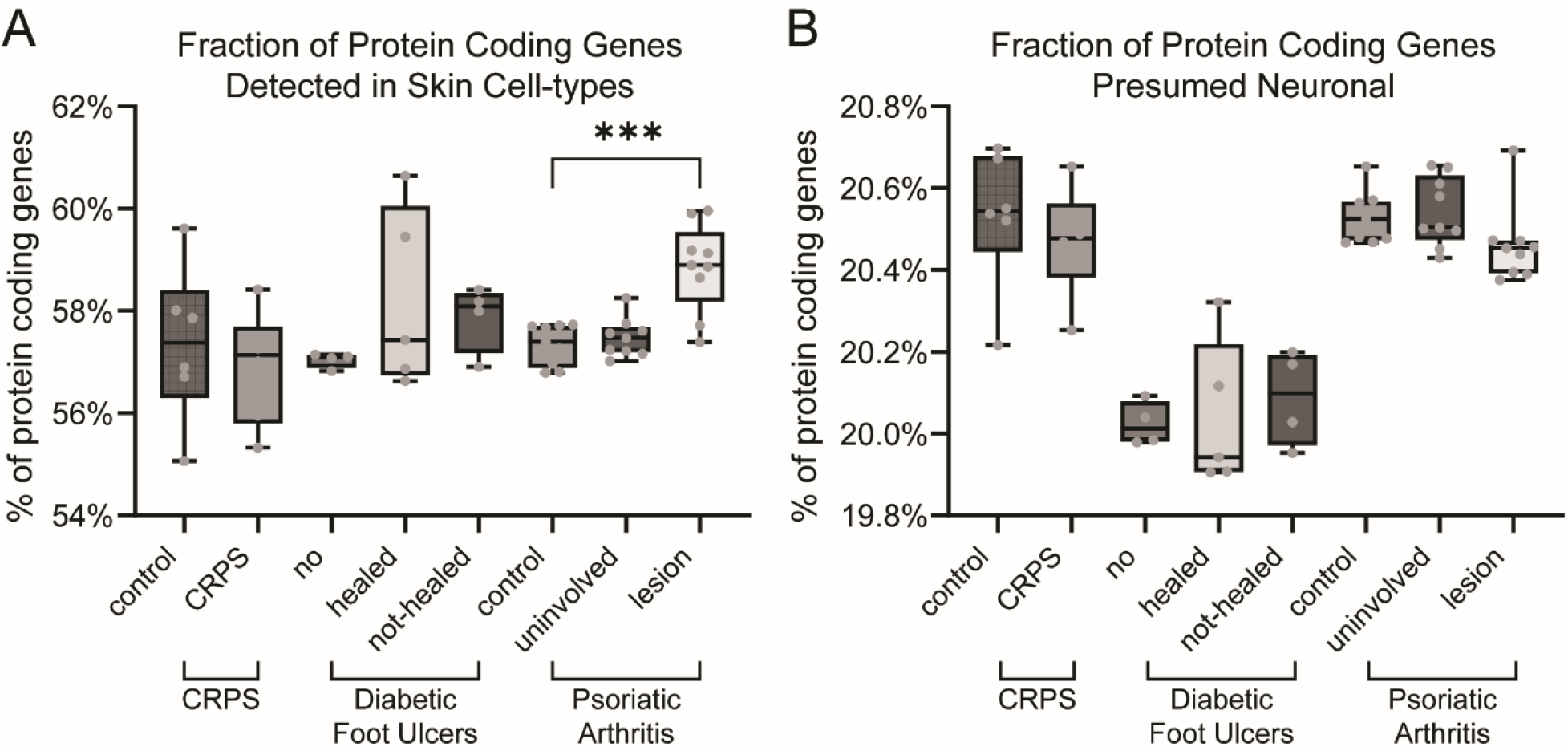
Validation of neuronal gene detection in other skin bulk RNA-seq datasets. Fraction of protein coding genes detected in bulk RNA-seq data of skin biopsies that are A) detected in cell-types identified in skin scRNA-seq datasets B) presumed expressed by nerve endings in the skin. *** p-value < 0.00

Having established direct spatial evidence for neuronal transcripts in human skin, we next asked whether this analytical framework could still detect presumed neuronal gene expression in other neuropathic conditions. We examined whether neuron-specific genes could be detected in skin biopsy samples from additional painful neuropathies known to involve epidermal nerve fiber loss [27; 47], including 12 limb biopsies from a complex regional pain syndrome (CRPS) study (paired affected and unaffected sites from 6 CRPS patients) [18], 13 volar forearm biopsies from a diabetic painful neuropathy (DPN) study (all diabetic patients: 5 with healed ulcers, 4 with non-healed ulcers, and 4 with no ulcer) [44], and 27 biopsies from a psoriatic arthritis study (9 lesional, 9 non-lesional, and 9 healthy-control samples) [9]. Across all three diseases, the number of expressed protein-coding genes from skin cell types (based on the skin scRNA-seq reference) did not differ significantly between disease and control conditions, with the exception of lesional samples from the psoriatic arthritis dataset, which is consistent with the acute inflammation and altered keratinization that occur at lesion sites [11]. Notably, within-dataset comparisons showed no significant differences in the detection of presumed neuronal genes across any of the three studies. This indicates that, despite the widespread occurrence of nerve-ending die-back in these neuropathic skin diseases, the proposed approach reliably profiles neuronal gene expression in skin biopsy samples even under severe conditions such as ulceration or active lesions. It is important to note that these biopsies are not specific for the epidermis and still contain deeper skin layers where nerve fibers may still be abundant even in patients with intraepidermal nerve fiber loss.

## Discussion

A major issue in understanding mechanisms of neuropathic pain is our inability to sample the nervous system in most patients. Great strides have been made in understanding neuropathic pain in humans by using DRGs or nerve samples obtained post-mortem [14; 35; 42], or during rare surgeries [1; 17; 26; 29; 32; 41; 46], but this still accounts for only a tiny portion of neuropathic pain patients, and these kinds of samples cannot, for the most part, be used to understand pain mechanisms in individual patients or to make treatment decisions. Skin biopsies, on the other hand, can easily be obtained from almost any neuropathic pain patient, can be obtained at multiple time points, and their assessment can be incorporated into treatment decisions and/or used in clinical trials. We went into this study with a simple hypothesis that skin biopsies could be used to assess the expression of DRG neuronal mRNAs in a neuropathic pain cohort. Our work strongly supports the utility of this approach and warrants more extensive, prospectively designed studies to better understand how skin biopsies can be used to understand sensory neuron pathology at the transcriptional level in neuropathic pain patients.

While we could confidently call expression of most DEGs found in the skin biopsies of these PIPN patients in either skin cells or DRG neurons, there were still 227 DEGs not detected in either scRNA-seq datasets from skin samples or transcriptomic datasets from human DRG neurons [1; 4; 42]. This is likely due to the limitation of sc/snRNA-seq sequencing depth. In addition, certain library-preparation chemistries, such as 10x Genomics Flex, capture only a specific set of protein-coding transcripts.

These approaches miss certain less studied or newly annotated genes, like KCNA6, which likely contribute to substantial fraction of the remaining undetected genes. In this study, the used skin reference dataset integrated 273,178 cells from 15 anatomical regions across 85 individuals. Across these samples, 9,533 genes were detected in at least one cell type, whereas 18,497 genes were identified in bulk skin RNA-seq datasets, underscoring the reduced sensitivity of single-cell approaches. Accordingly, it is plausible that some of the 227 differentially expressed genes (DEGs) not detected in the scRNA-seq references are nonetheless expressed in skin cells at levels below the detection threshold. Notably, the 639 neuronal genes inferred to originate from nerve terminals likely represent a conservative estimate, as ongoing nerve-ending degeneration may further limit their detection. Importantly, the primary objective of this study was to demonstrate the feasibility of detecting nerve terminal signatures in human skin and to use these data to infer subsets of sensory neurons present in the tissue; although we acknowledge that current technical limitations preclude comprehensive characterization of these nerve endings, such exhaustive profiling can be done in the future with advanced spatial transcriptomics techniques and improved cell segmentation technologies.

Another possibility, which we favor, is that our inability to detect many of these 227 DEGs reflects changes in gene expression in skin cells in PIPN. Here it is notable that both animal studies and human experiments in PIPN patients support the conclusion that certain subtypes of skin cells show physiological alterations in PIPN. There is a growing literature on changes in keratinocytes in PIPN and CIPN, with these cells likely playing an important role in cell-neuron communication in the skin that is involved in promoting aspects of neuropathic pain both in animal CIPN models and in human patients. From this perspective it is notable that among the upregulated DEGs that were expressed in skin cells from the control scRNA-seq datasets we used, most of the genes that showed enriched expression in a particular cell type were expressed by cycling keratinocytes. We think it is likely that future single cell sequencing experiments on skin biopsies from PIPN patients will reveal dysregulated gene expression in keratinocytes that may correlate with pain severity.

An additional limitation concerns sex balance: all 6 PIPN samples analyzed here were obtained from female patients. Several of the reference datasets are also not balanced in sex — for example, the CRPS dataset is predominantly female (reflecting the higher prevalence of CRPS in women), whereas the foreskin scRNA-seq data are exclusively male. We acknowledge this limitation and plan to incorporate more balanced cohorts in future work to systematically evaluate the effect of CIPN on skin nerve endings in skin biopsy samples.

We also note that, although nerve-ending die-back has been widely reported across many neuropathic conditions, the PIPN samples analyzed here did not show significant reductions in IENF density, despite CIPN20 scores and other clinical assessments indicating painful neuropathy. Comparable numbers of presumed nerve-ending-derived genes were detected in skin biopsies from three additional neuropathy-associated diseases in which die-back has been documented, suggesting that this approach is robust across multiple clinical settings. Detection may be robust for neuronal genes because deeper axons are intact and present in these skin biopsies.

Further work using spatial transcriptomics can be done to localize these RNAs more specifically to axons in different locations in the skin. Our Xenium results demonstrate that this approach is technically feasible, although there is an obvious tradeoff in cost for spatial transcriptomics versus bulk RNA-seq for potential clinical application.

Our analysis of skin bulk RNA-seq data revealed that many DEGs originated from neuronal populations, highlighting neuron-specific changes in the skin of PIPN patients. As noted above, we concede that it will be important to evaluate changes in gene expression in using single cell sequencing in PIPN patient skin biopsies so these changes in gene expression can be definitively assigned to neurons. This can also be done through the use of spatial transcriptomic technologies like 10X Genomics Xenium that has the resolution to detect axonal transcripts, as we have already done here. We think it is likely that such studies will confirm that the DEGs we detected in this study can be assigned to neuronal expression for two reasons. First, many of these genes encode ion channels that are already known to be expressed primarily in neurons and other excitable cells that do not have cell bodies that are present in the skin. Second, the observation that many down-regulated DEGs that were expressed in neurons but not skin cells could be assigned to the *PDIA2* population is consistent with die-back of certain classes of injured axons in PIPN.

In conclusion, our study provides support for the detection of neuronal mRNAs, localized to axons and/or nerve endings, among DEGs in bulk RNA sequencing experiments from patients with PIPN and other types of peripheral neuropathy. We propose that relatively inexpensive bulk RNA-seq of skin biopsies from patients with neuropathic pain offers valuable insights into disease mechanisms. This technique is readily adaptable for future research initiatives and shows promise as a novel clinical diagnostic tool to inform precision medicine approaches. This technique has been used on DRGs [1; 26; 28] and nerves [32; 41] obtained from rare surgeries to gain important mechanistic insight into neuropathic pain but these tissues are inaccessible in most patients. The findings described here demonstrate that it is possible to use skin biopsies, which are commonly obtained in routine clinical practice to measure differential gene expression in the nervous endings of neuropathic pain patients.

## Acknowledgements

This research was supported by the National Institute of Neurological Disorders and Stroke of the National Institutes of Health through the PRECISION Human Pain Network (RRID:SCR_025458), part of the NIH HEAL Initiative (https://heal.nih.gov/) under award number U19NS130608 to TJP. This research was supported by the National Cancer Institute of the National Institutes of Health via Award 1R01CA215973 to SR and R01CA275870 to NPS. The content is solely the responsibility of the authors and does not necessarily represent the official views of the National Institutes of Health.

## References

[1] Arendt-Tranholm A, Sankaranarayanan I, Payne C, Moreno MM, Mazhar K, Yap N, Chiu AP, Barry A, Patel PP, Inturi NN, Ferreira DT, Amin A, Karandikar M, Jarvik JG, Turner JA, Hofstetter CP, Curatolo M, Price TJ. Single-cell characterization of the human C2 dorsal root ganglion recovered from C1-2 arthrodesis surgery: implications for neck pain. Brain 2025(in press).

[2] Baba Y, Maezawa Y, Kondo N, Minamizuka T, Udagawa H, Ide S, Ide K, Teramoto N, Yamaguchi A, Kaneko H, Funayama S, Kato H, Shoji M, Aono K, Miyabayashi M, Sato T, Kitamoto T, Yagyu Y, Ishibashi R, Koshizaka M, Endo Y, Kanda M, Takemoto M, Takayama N, Yasuda K, Kobayashi Y, Yokote K. Tcf21 modulates fibroblast activation and promotes cardiac fibrosis after injury via Pdgfrb signaling. Sci Rep 2025;15(1):28260.

[3] Bhuiyan SA, Nagi SS, Sankaranarayanan I, Semizoglou E, Usoskin D, Yang L, Yu H, Arendt-Tranholm A, Bertels Z, Bhatia P, Bouchatta O, Boyer K, Cervantes A, Chalif J, Chintalapudi H, Cicalo A, Copits B, Cronin C, Curatolo M, Dong X, Dougherty PM, Dourson A, Funk G, Gabriel K, Griesemer DS, Guo H, Gupta P, Hofstetter C, Horton P, Hsieh A, Inturi NN, Jain A, Jayakar S, Johnston B, Kim R, Krauter D, Kupari J, Lemen J, Lesnak JB, Liu W, Lopez I, Lu Y, MacMillan HJ, Mazhar K, Meriau P, Moffitt JR, Moreno MM, Mwirigi JM, Naz H, O’Brein J, Payne M, Rosario JD, Rosen SF, Shiers S, Simpson E, Slivicki R, Stone JR, Tavares-Ferreira D, Uhelski M, Woolf CJ, Xu Q, Yi J, Yousuf MS, Zhu D, Cavalli V, Zhao G, Olausson H, Ernfors P, Gereau RW, Luo W, Price TJ, Renthal W, Network NPHP. A Reference Atlas of the Human Dorsal Root Ganglion. bioRxiv 2025–11–06.

[4] Bhuiyan SA, Xu M, Yang L, Semizoglou E, Bhatia P, Pantaleo KI, Tochitsky I, Jain A, Erdogan B, Blair S, Cat V, Mwirigi JM, Sankaranarayanan I, Tavares-Ferreira D, Green U, McIlvried LA, Copits BA, Bertels Z, Del Rosario JS, Widman AJ, Slivicki RA, Yi J, Sharif-Naeini R, Woolf CJ, Lennerz JK, Whited JL, Price TJ, Renthal W. Harmonized cross-species cell atlases of trigeminal and dorsal root ganglia. Science Advances 2024;10(25):eadj9173.

[5] Devigili G, Rinaldo S, Lombardi R, Cazzato D, Marchi M, Salvi E, Eleopra R, Lauria G. Diagnostic criteria for small fibre neuropathy in clinical practice and research. Brain 2019;142(12):3728–3736.

[6] Duan HX, Li BW, Zhuang X, Wang LT, Cao Q, Tan LH, Qu GF, Xiao S. TCF21 inhibits tumor-associated angiogenesis and suppresses the growth of cholangiocarcinoma by targeting PI3K/Akt and ERK signaling. Am J Physiol Gastrointest Liver Physiol 2019;316(6):G763–G773.

[7] Engelstad JK, Taylor SW, Witt LV, Hoebing BJ, Herrmann DN, Dyck PJ, Klein CJ, Johnson DM, Davies JL, Carter RE, Dyck PJ. Epidermal nerve fibers: confidence intervals and continuous measures with nerve conduction. Neurology 2012;79(22):2187–2193.

[8] Giorgi S, Lamberti A, Butron L, Gross-Amat O, Alarcon-Alarcon D, Rodriguez-Canas E, Fernandez-Carvajal A, Ferrer-Montiel A. Compartmentalized primary cultures of dorsal root ganglion neurons to model peripheral pathophysiological conditions.Mol Pain 2023;19:17448069231197102.

[9] H J, J C, IB M, G G, S S. Differences in transcriptional changes in psoriasis and psoriatic arthritis skin with immunoglobulin gene enrichment in psoriatic arthritis - PubMed. Rheumatology (Oxford, England) 01/04/2024;63(1).

[10] Jimenez-Diaz L, Geranton SM, Passmore GM, Leith JL, Fisher AS, Berliocchi L, Sivasubramaniam AK, Sheasby A, Lumb BM, Hunt SP. Local translation in primary afferent fibers regulates nociception. PLoS One 2008;3(4):e1961.

[11] Johnsson H, Cole J, Siebert S, McInnes IB, Graham G. Cutaneous lesions in psoriatic arthritis are enriched in chemokine transcriptomic pathways. Arthritis Research & Therapy 2023 May 2;25(1).

[12] Khoutorsky A, Price TJ. Translational Control Mechanisms in Persistent Pain. Trends Neurosci 2018;41(2):100–114.

[13] Kuleshov MV, Jones MR, Rouillard AD, Fernandez NF, Duan Q, Wang Z, Koplev S, Jenkins SL, Jagodnik KM, Lachmann A, McDermott MG, Monteiro CD, Gundersen GW, Ma’ayan A. Enrichr: a comprehensive gene set enrichment analysis web server 2016 update. Nucleic Acids Res 2016;44(W1):W90–97.

[14] Lacagnina MJ, Willcox KF, Boukelmoune N, Bavencoffe A, Sankaranarayanan I, Barratt DT, Zuberi YA, Dayani D, Chavez MV, Lu JT, Farinotti AB, Shiers S, Barry AM, Mwirigi JM, Tavares-Ferreira D, Funk GA, Cervantes AM, Svensson CI, Walters ET, Hutchinson MR, Heijnen CJ, Price TJ, Fiore NT, Grace PM. B cells drive neuropathic pain-related behaviors in mice through IgG-Fc gamma receptor signaling. Sci Transl Med 2024;16(766):eadj1277.

[15] Li Y, Adamek P, Zhang H, Tatsui CE, Rhines LD, Mrozkova P, Li Q, Kosturakis AK, Cassidy RM, Harrison DS, Cata JP, Sapire K, Zhang H, Kennamer-Chapman RM, Jawad AB, Ghetti A, Yan J, Palecek J, Dougherty PM. The Cancer Chemotherapeutic Paclitaxel Increases Human and Rodent Sensory Neuron Responses to TRPV1 by Activation of TLR4. Journal of Neuroscience 2015–09–30;35(39).

[16] Li Y, North RY, Rhines LD, Tatsui CE, Rao G, Edwards DD, Cassidy RM, Harrison DS, Johansson CA, Zhang H, Dougherty PM. DRG Voltage-Gated Sodium Channel 1.7 Is Upregulated in Paclitaxel-Induced Neuropathy in Rats and in Humans with Neuropathic Pain. J Neurosci 2018;38(5):1124–1136.

[17] Li Y, Uhelski ML, North RY, Mwirigi JM, Tatsui CE, McDonough KE, Cata JP, Corrales G, Dussor G, Price TJ, Dougherty PM. Tomivosertib reduces ectopic activity in dorsal root ganglion neurons from patients with radiculopathy. Brain 2024;147(9):2991–2997.

[18] M PVV, A J, P B, J D, H R, L O, L S, P S, S C, O D, F B, S D. Identification of two biological subgroups of complex regional pain syndrome type 1 by transcriptomic profiling of skin and blood in women - PubMed. Molecular medicine (Cambridge, Mass) 03/12/2025;31(1).

[19] Ma J, Kavelaars A, Dougherty PM, Heijnen CJ. Beyond symptomatic relief for chemotherapy-induced peripheral neuropathy: Targeting the source. Cancer 2018;124(11):2289–2298.

[20] Megat S, Ray PR, Moy JK, Lou TF, Barragan-Iglesias P, Li Y, Pradhan G, Wanghzou A, Ahmad A, Burton MD, North RY, Dougherty PM, Khoutorsky A, Sonenberg N, Webster KR, Dussor G, Campbell ZT, Price TJ. Nociceptor Translational Profiling Reveals the Ragulator-Rag GTPase Complex as a Critical Generator of Neuropathic Pain. J Neurosci 2019;39(3):393–411.

[21] Melemedjian OK, Asiedu MN, Tillu DV, Peebles KA, Yan J, Ertz N, Dussor GO, Price TJ. IL-6-and NGF-Induced Rapid Control of Protein Synthesis and Nociceptive Plasticity via Convergent Signaling to the eIF4F Complex. J Neurosci 2010;30(45):15113–15123.

[22] Melemedjian OK, Tillu DV, Moy JK, Asiedu MN, Mandell EK, Ghosh S, Dussor G, Price TJ. Local translation and retrograde axonal transport of CREB regulates IL-6-induced nociceptive plasticity. Mol Pain 2014;10:45.

[23] Mikesell AR, Isaeva E, Schulte ML, Menzel AD, Sriram A, Prahl MM, Shin SM, Sadler KE, Yu H, Stucky CL. Keratinocyte Piezo1 drives paclitaxel-induced mechanical hypersensitivity. bioRxiv 2023.

[24] Mikesell AR, Isaeva E, Schulte ML, Menzel AD, Sriram A, Prahl MM, Shin SM, Sadler KE, Yu H, Stucky CL. Increased keratinocyte activity and PIEZO1 signaling contribute to paclitaxel-induced mechanical hypersensitivity. Sci Transl Med 2024;16(777):eadn5629.

[25] Mohr E, Richter D. Axonal mRNAs: functional significance in vertebrates and invertebrates. J Neurocytol 2000;29(11-12):783–791.

[26] North RY, Li Y, Ray P, Rhines LD, Tatsui CE, Rao G, Johansson CA, Zhang H, Kim YH, Zhang B, Dussor G, Kim TH, Price TJ, Dougherty PM. Electrophysiological and transcriptomic correlates of neuropathic pain in human dorsal root ganglion neurons. Brain 2019;142(5):1215–1226.

[27] Peltier AC, Myers MI, Artibee KJ, Hamilton AD, Yan Q, Guo J, Shi Y, Wang L, Li J. Evaluation of dermal myelinated nerve fibers in diabetes mellitus. Journal of the peripheral nervous system : JPNS 2013 Jun;18(2).

[28] Ray PR, Shiers S, Caruso JP, Tavares-Ferreira D, Sankaranarayanan I, Uhelski ML, Li Y, North RY, Tatsui C, Dussor G, Burton MD, Dougherty PM, Price TJ. RNA profiling of human dorsal root ganglia reveals sex differences in mechanisms promoting neuropathic pain. Brain 2023;146(2):749–766.

[29] Ray PR, Shiers S, Tavares-Ferreira D, Sankaranarayanan I, Uhelski ML, Li Y, North RY, Tatsui C, Dussor G, Burton MD, Dougherty PM, Price TJ. RNA Profiling of Neuropathic Pain-Associated Human DRGs Reveal Sex-differences in Neuroimmune Interactions Promoting Pain. bioRxiv 2021:2021.2011.2027.470190.

[30] Restrepo P, Wilder A, Houser A, Sandhu HS, Ramirez A, Grace Hren M, Gill R, Kazmi A, Chen L, Nigro A, Imanishi I, Demircioglu D, Hasson D, Soto A, McQuillan S, Gonzalez-Kozlova E, Brody R, Ungar B, Kasper M, Lu CP, Torina P, Lewin JM, Gnjatic S, Ma S, Ji AL, Restrepo P, Wilder A, Houser A, Sandhu HS, Ramirez A, Grace Hren M, Gill R, Kazmi A, Chen L, Nigro A, Imanishi I, Demircioglu D, Hasson D, Soto A, McQuillan S, Gonzalez-Kozlova E, Brody R, Ungar B, Kasper M, Lu CP, Torina P, Lewin JM, Gnjatic S, Ma S, Ji AL. Single-cell spatial transcriptomic analysis of human skin anatomy. Nature Genetics 2026 58:4 2026–03–23;58(4).

[31] Ruangsri S, Lin A, Mulpuri Y, Lee K, Spigelman I, Nishimura I. Relationship of axonal voltage-gated sodium channel 1.8 (NaV1.8) mRNA accumulation to sciatic nerve injury-induced painful neuropathy in rats. J Biol Chem 2011;286(46):39836–39847.

[32] Sandy-Hindmarch OP, Chang PS, Scheuren PS, De Schoenmacker I, Hubli M, Loizou C, Wirth S, Mahadevan D, Wiberg A, Furniss D, Calvo M, Bennett DLH, Denk F, Baskozos G, Schmid AB. The local molecular signature of human peripheral neuropathic pain. Pain 2024.

[33] Seretny M, Currie GL, Sena ES, Ramnarine S, Grant R, MacLeod MR, Colvin LA, Fallon M. Incidence, prevalence, and predictors of chemotherapy-induced peripheral neuropathy: A systematic review and meta-analysis. Pain 2014;155(12):2461–2470.

[34] Shah A, Hoffman EM, Mauermann ML, Loprinzi CL, Windebank AJ, Klein CJ, Staff NP. Incidence and disease burden of chemotherapy-induced peripheral neuropathy in a population-based cohort. J Neurol Neurosurg Psychiatry 2018;89(6):636–641.

[35] Shiers SI, Mazhar K, Wangzhou A, Haberberger R, Lesnak JB, Ezeji NA, Sankaranarayanan I, Tavares-Ferreira D, Cervantes A, Funk G, Horton P, Vines E, Dussor G, Price TJ. Nageotte nodules in human dorsal root ganglia reveal neurodegeneration in diabetic peripheral neuropathy. Nat Commun 2025;16(1):4168.

[36] Staff NP, Fehrenbacher JC, Caillaud M, Damaj MI, Segal RA, Rieger S. Pathogenesis of paclitaxel-induced peripheral neuropathy: A current review of in vitro and in vivo findings using rodent and human model systems. Exp Neurol 2020;324:113121.

[37] Staff NP, Grisold A, Grisold W, Windebank AJ. Chemotherapy-induced peripheral neuropathy: A current review. Ann Neurol 2017;81(6):772–781.

[38] Staff NP, Hrstka SC, Dasari S, Capobianco E, Rieger S. Skin Extracellular Matrix Breakdown Following Paclitaxel Therapy in Patients with Chemotherapy-Induced Peripheral Neuropathy. Cancers (Basel) 2023;15(16).

[39] Stucky CL, Mikesell AR. Cutaneous pain in disorders affecting peripheral nerves. Neurosci Lett 2021;765:136233.

[40] Tavares-Ferreira D, Shen BQ, Mwirigi JM, Shiers S, Sankaranarayanan I, Sreerangapuri A, Kotamarti MB, Inturi NN, Mazhar K, Ubogu EE, Thomas GL, Lalli T, Rozen SM, Wukich DK, Price TJ. Cell and molecular profiles in peripheral nerves shift toward inflammatory phenotypes in diabetic peripheral neuropathy. J Clin Invest 2025;135(20).

[41] Tavares-Ferreira D, Shen BQ, Mwirigi JM, Shiers S, Sankaranarayanan I, Sreerangapuri A, Kotamarti MB, Inturi NN, Mazhar K, Ubogu EE, Thomas GL, Lalli T, Rozen SM, Wukich DK, Price TJ. Cell and molecular profiles in peripheral nerves shift toward inflammatory phenotypes in diabetic peripheral neuropathy. J Clin Invest 2025.

[42] Tavares-Ferreira D, Shiers S, Ray PR, Wangzhou A, Jeevakumar V, Sankaranarayanan I, Cervantes AM, Reese JC, Chamessian A, Copits BA, Dougherty PM, Gereau RWt, Burton MD, Dussor G, Price TJ. Spatial transcriptomics of dorsal root ganglia identifies molecular signatures of human nociceptors. Sci Transl Med 2022;14(632):eabj8186.

[43] Thakor DK, Lin A, Matsuka Y, Meyer EM, Ruangsri S, Nishimura I, Spigelman I. Increased peripheral nerve excitability and local NaV1.8 mRNA up-regulation in painful neuropathy. Mol Pain 2009;5:14.

[44] Theocharidis G, Baltzis D, Roustit M, Tellechea A, Dangwal S, Khetani RS, Shu B, Zhao W, Fu J, Bhasin S, Kafanas A, Hui D, Sui SH, Patsopoulos NA, Bhasin M, Veves A. Integrated Skin Transcriptomics and Serum Multiplex Assays Reveal Novel Mechanisms of Wound Healing in Diabetic Foot Ulcers. Diabetes 2020;69(10):2157–2169.

[45] Trecarichi A, Flatters SJL. Mitochondrial dysfunction in the pathogenesis of chemotherapy-induced peripheral neuropathy. Int Rev Neurobiol 2019;145:83–126.

[46] Uhelski ML, Schaub MK, Espinosa F, Heles M, Cortes N, Li Y, Tatsui CE, Rhines LD, North RY, Alvarez-Breckendridge C, Funk G, Horton P, Cervantes A, Lesnak JB, Curalto M, Dussor G, Price TJ, Dougherty PM. Suzetrigine (VX-548) exhibits activity-dependent effects on human dorsal root ganglion neurons. bioRxiv 2025:2025.2004.2009.648014.

[47] VF R, P K, PD D, EL S, AJ T, TS J, LF K. Bilaterally Reduced Intraepidermal Nerve Fiber Density in Unilateral CRPS-I-PubMed. Pain medicine (Malden, Mass) 10/01/2018;19(10).

[48] Xie Z, Bailey A, Kuleshov MV, Clarke DJB, Evangelista JE, Jenkins SL, Lachmann A, Wojciechowicz ML, Kropiwnicki E, Jagodnik KM, Jeon M, Ma’ayan A. Gene Set Knowledge Discovery with Enrichr. Curr Protoc 2021;1(3):e90.

[49] Zhang H, Dougherty PM. Enhanced Excitability of Primary Sensory Neurons and Altered Gene Expression of Neuronal Ion Channels in Dorsal Root Ganglion in Paclitaxel-induced Peripheral Neuropathy. Anesthesiology June 2014;120(6).

