## Supplemental Table 1 for "Detection of probable neuronal gene expression changes in skin biopsies from patients with paclitaxel-induced peripheral neuropathy"

METHODS: After IRB approval, three female breast cancer patients with neuropathy following weekly paclitaxel chemotherapy and three matched healthy controls (ages 60–70; **Figure**) were enrolled as described previously (PMID 37627219).

Neurological history and examination, Neuropathy Impairment Scores in the lower limbs (NIS-LL), and QLQ-CIPN20 quality of life questionnaires were administered to all participants by a neurologist with peripheral nerve fellowship training (NPS). The NIS-LL provides a composite assessment of strength, sensory loss, and diminished deep tendon reflexes in the lower extremities (PMID: 9222195). The QLQ-CIPN20, a validated instrument for detecting CIPN (PMID: 15911236), consists of 20 items evaluating multiple aspects of CIPN-related symptoms and their effects on daily functioning and quality of life. This tool encompasses diverse domains, including sensory symptoms (e.g., tingling, numbness, pain), motor symptoms (e.g., weakness, coordination difficulties), autonomic symptoms (e.g., abnormal sweating, gastrointestinal disturbances), and functional impairments (e.g., challenges with fine motor skills, ambulation, and activities of daily living). Participants rated the severity and frequency of these symptoms over the previous week using a scale from 1 (not at all) to 4 (very much). Additionally, the questionnaire includes items addressing participants' perceptions of the overall impact of CIPN on physical, emotional, and social well-being. Collectively, these measures offer valuable insight into subjective experiences and facilitate quantification of the burden of CIPN symptoms on everyday life.

Following informed consent, two 3mm skin punch biopsies were performed 10 cm proximal to the lateral malleolus on the distal leg, as described previously (PMID 23100396). The biopsies were carried out following local injection of 2% lidocaine with epinephrine anesthesia under sterile conditions. Skin biopsies were processed for PGP9.5 staining, a marker for sensory nerve endings in the epidermis, and quantification of ENFD were performed at the Mayo Clinic Peripheral Nerve Laboratory according to standard clinical practice procedures (regulated by Clinical Laboratory Improvements Amendments—CLIA).

RESULTS:

Clinical Descriptions: Three female participants between 60 and 70 years of age with invasive breast carcinoma received adjuvant paclitaxel therapy (80 mg/m²/week for 12 weeks), during which they subsequently experienced neuropathic symptoms (**Figure**). Participants in this study were recruited to assess neuropathic symptoms, determine epidermal nerve fiber density (ENFD), and examine skin biology related to CIPN. None of the participants had previous neuropathic symptoms, a family history of neuropathy, or prior chemotherapy before receiving the paclitaxel regimen described. Subject 1 (CIPN001), a 70-year-old with invasive lobular carcinoma, developed numbness, coldness, tingling, and burning sensations predominantly in the toes during the final treatment cycle. Enrolled 35 weeks post-therapy, she recorded an EORTC QLQ-CIPN20 score of 31, NIS-LL score of 2, and ENFD mean of 8.9 (95% CI 7.5–10.4; estimated 5th percentile 2.5). Subject 2 (CIPN002), also 70 years old with invasive lobular carcinoma, reported tingling, numbness, and burning in her fingers and feet beginning in the second cycle, which necessitated a 25% paclitaxel dose reduction for the final six cycles. She was enrolled 31 weeks after her last paclitaxel administration with a QLQ-CIPN20 score of 34, NIS-LL score of 4, and ENFD mean of 5.3 (95% CI 4.3–6.6; estimated 5th percentile 2.5). Subject 3 (CIPN003), aged 60 with invasive ductal carcinoma, developed tingling and numbness during the fifth cycle. She joined the study five weeks following her last paclitaxel dose, reporting a QLQ-CIPN20 score of 25, NIS-LL score of 2, and ENFD mean of 9.6 (95% CI 8.2–11.2; estimated 5th percentile 3.1). All subjects demonstrated normal muscle strength on neurological examination, with varying levels of sensory impairment and diminished deep tendon reflexes. Electrodiagnostic assessment was not performed as part of clinical evaluation or study protocol.  Three age- and sex-matched control subjects (aged 64, 65, and 70) were recruited, and determined to not have peripheral neuropathy. Mean ENFD for the control subjects were 14.9, 7.6, and 7.6, respectively.  All recruited subjects had ENFD that was normal for age and sex, and there was no statistical significance in ENFD between CIPN and control subjects ($p>0.05$).

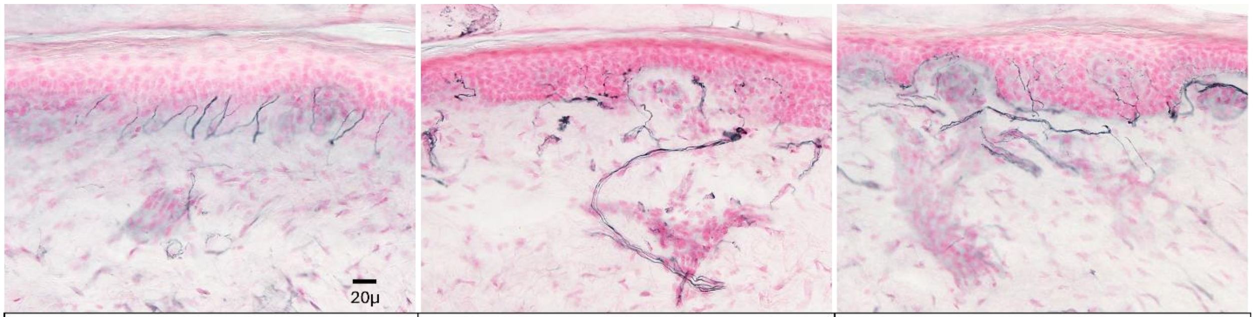


| CIPN001 | CIPN002 | CIPN003 |
|---|---|---|
| **AGE:** 70 | **AGE:** 70 | **AGE:** 60 |
| **SEX:** F | **SEX:** F | **SEX:** F |
| **CANCER:** invasive lobular breast carcinoma | **CANCER:** invasive lobular breast carcinoma | **CANCER:** invasive ductal breast carcinoma |
| **PACLITAXEL:** 80 mg/m$^2$/week for 12 weeks | **PACLITAXEL:** 80 mg/m$^2$/week for 6 weeks, 60 mg/m$^2$/week for 6 weeks | **PACLITAXEL:** 80 mg/m$^2$/week for 12 weeks |
| **WEEKS FROM CIPN TO BIOPSY:** 35 | **WEEKS FROM CIPN TO BIOPSY:** 31 | **WEEKS FROM CIPN TO BIOPSY:** 5 |
| **NIS-LL**: 2 | **NIS-LL**: 12 | **NIS-LL**: 2 |
| **QLQ-CIPN20 (total):** 31 | **QLQ-CIPN20 (total):** 34 | **QLQ-CIPN20 (total):** 25 |
| **QLQ-CIPN20 (sensory):** 22 | **QLQ-CIPN20 (sensory):** 22 | **QLQ-CIPN20 (sensory):** 16 |
| **QLQ-CIPN20 (motor):** 3 | **QLQ-CIPN20 (motor):** 4 | **QLQ-CIPN20 (motor):** 3 |
| **ENFD:** *mean*: 8.9 (95% *CI* 7.5-10.4), *5$^{th}$ percentile for age & sex*: 2.5 | **ENFD:** *mean*: 5.3 (95% *CI* 4.3-6.6), *5$^{th}$ percentile for age & sex*: 2.5 | **ENFD:** *mean*: 9.6 (95% *CI* 8.2-11.2), *5$^{th}$ percentile for age & sex*: 3.1 |

FIGURE LEGEND: Clinical Data from three subjects with paclitaxel-induced peripheral neuropathy. Photomicrographs of skin immunostained with antibodies to PGP9.5 counterstained with Fast Red.

ACKNOWLEDGEMENTS: We thank JaNean Engelstad and Peter Dyck in the Mayo Clinic Peripheral Nerve Laboratory for their valuable assistance with IENF quantification.
